# Genomic Locus Modulating Corneal Thickness in the Mouse Identifies *POU6F2* as a Potential Risk of Developing Glaucoma

**DOI:** 10.1101/212605

**Authors:** Rebecca King, Felix L. Struebing, Ying Li, Jiaxing Wang, Allison Ashley Koch, Jessica Cooke Bailey, Puya Gharahkhani, International Glaucoma Genetics Consortium, NEIGHBORHOOD consortium, Stuart MacGregor, R. Rand Allingham, Michael A. Hauser, Janey L. Wiggs, Eldon E. Geisert

**Affiliations:** Department of Ophthalmology, Emory University, Atlanta, Georgia United States of America; Department of Ophthalmology, Tianjin Medical University General Hospital, Tianjin, China.; Center for Human Disease Modeling, Duke University, Durham, North Carolina, United States of America; Department of Epidemiology and Biostatistics, Case Western Reserve University, Cleveland, Ohio, United States of America; Statistical Genetics, QIMR Berghofer Medical Research Institute, Brisbane, Queensland, Australia; Department of Medicine and Ophthalmology, Duke University Medical Center Durham, North Carolina, United States of America; Department of Ophthalmology, Harvard Medical School of Medicine, Massachusetts Eye and Ear Infirmary, Boston, Massachusetts, United States of America

**Author notes:** Corresponding Author: Eldon E. Geisert E-Mail (EEG). Author Contribution Statement: Rebecca King maintained the animal colonies and collected all of the CCT data at Emory University. She also had a primary role in the analysis of the entire set of BXD CCT data and contributed all of the immunohistochemistry to the study. Felix L. Struebing was involved in the analysis of the BXD mouse data and the PCR analysis of gene expression in the mouse retina and mouse cornea. He was responsible for the immunostaining of the adult retina. Ying Li stained and quantified the flat mounted retinas to determine the number and POU6F2-positive ganglion cells. She was involved in validating the antibody used to stain for POU6F2. Jiaxing Wang conducted all of the experiments examining the validity of probe sets on the Affymetrix Chip. He examined the distribution of *Pou6f2* in different tissues using PCR. Jessica Cooke Bailey assisted with the analyses using the NEIGHBORHOOD glaucoma dataset. Michael A. Hauser designed and implemented the analysis of the human CCT data. Allison Ashley Koch designed and implemented the analysis of the human CCT data. R. Rand Allingham designed and implemented the analysis of the human CCT data. Stuart MacGregor directed the analysis of human CCT within the International Glaucoma Genetics Consortium. Puya Gharahkhani assisted with the analysis of the human CCT data. Janey L. Wiggs was involved in the initial design of the experiments and analysis of the NEIGHBORHOOD glaucoma dataset. Eldon E. Geisert had direct oversight of the BXD mouse study, designed the experiments and wrote the manuscript.

## Abstract

Purpose: Central corneal thickness (CCT) is one of the most heritable ocular traits and it is also a phenotypic risk factor for primary open angle glaucoma (POAG). The present study uses the BXD Recombinant Inbred (RI) strains to identify novel quantitative trait loci (QTLs) modulating CCT in the mouse with the potential of identifying a molecular link between CCT and risk of developing POAG.

Methods: The BXD RI strain set was used to define mammalian genomic loci modulating CCT, with a total of 818 corneas measured from 61 BXD RI strains (between 60-100 days of age). The mice were anesthetized and the eyes were positioned in front of the lens of the Phoenix Micron IV Image-Guided OCT system or the Bioptigen OCT system. CCT data for each strain was averaged and used to identify quantitative trait loci (QTLs) modulating this phenotype using the bioinformatics tools on GeneNetwork (<u>www.genenetwork.org</u>). The candidate genes and genomic loci identified in the mouse were then directly compared with the summary data from a human primary open-angle glaucoma (POGA) genome wide association study (NEIGHBORHOOD) to determine if any genomic elements modulating mouse CCT are also risk factors for POAG.

Results: This analysis revealed one significant QTL on Chr 13 and a suggestive QTL on Chr 7. The significant locus on Chr 13 (13 to 19 Mb) was examined further to define candidate genes modulating this eye phenotype. For the Chr 13 QTL in the mouse, only one gene in the region (*Pou6f2*) contained nonsynonymous SNPs. Of these five nonsynonymous SNPs in *Pou6f2,* two resulted in changes in the amino acid proline which could result in altered secondary structure affecting protein function. The 7 Mb region under the mouse Chr 13 peak distributes over 2 chromosomes in the human: Chr 1 and Chr 7. These genomic loci were examined in the NEIGHBORHOOD database to determine if they are potential risk factors for human glaucoma identified using meta-data from human GWAS. The top 50 hits all resided within one gene (*POU6F2*), with the highest significance level of p = 10^−6^ for SNP rs76319873. POU6F2 is found in retinal ganglion cells and in corneal limbal stem cells. To test the effect of POU6F2 on CCT we examined the corneas of a *Pou6f2*-null mice and the corneas were thinner than those of wild-type littermates. In addition, these POU6F2 RGCs die early in the DBA/2J model of glaucoma than most RGCs.

Conclusions: Using a mouse genetic reference panel, we identified a transcription factor, *Pou6f2*, that modulates CCT in the mouse. POU6F2 is also found in a subset of retinal ganglion cells and these RGCs are sensitive to injury.

**Authors Summary:** Glaucoma is a complex group of diseases with several known causal mutations and many known risk factors. One well-known risk factor for developing primary open angle glaucoma is the thickness of the central cornea. The present study leverages a unique blend of systems biology methods using BXD recombinant inbred mice and genome-wide association studies from humans to define a putative molecular link between a phenotypic risk factor (central corneal thickness) and glaucoma. We identified a transcription factor, POU6F2, that is found in the developing retinal ganglion cells and cornea. POU6F2 is also present in a subpopulation of retinal ganglion cells and in stem cells of the cornea. Functional studies reveal that POU6F2 is associated the central corneal thickness and with susceptibility of retinal ganglion cells to injury.

## Introduction

Since the early Ocular Hypertension Treatment Studies (OHTS) [1] and subsequent independent findings of others [2, 3], central corneal thickness (CCT) was identified as a factor related to the risk for developing primary open angle glaucoma (POAG). In these studies, CCT was a powerful predictor for developing POAG, with thinner corneas being associated with an increased risk of developing POAG [1–3] and this risk was independent of the confounding effects of CCT on intraocular pressure measurements [1, 3]. The thinner CCT is also associated with an increased severity of visual field loss and a more rapid progression of the disease [4–6]. Furthermore, ethnic differences in CCT are correlated with increased risk of developing POAG and an increased severity of the disease[7]. Thus, there is a profound link between CCT and the risk of developing POAG.

In humans, there is a considerable variation in CCT, ranging from under 450 μm to over 650 μm [8–10] with a mean CCT of approximately 550 μm [8–13]. The variation in CCT is a highly heritable ocular trait, the genomic contribution of CCT is estimated to be near 90% [8, 9, 11, 14]. Two studies on monozygotic and dizygotic twins from two different human populations confirmed the heritability, with the Chinese population having a heritability of 0.88 [8] [9] and the population in Australia and the United Kingdom having a heritability of 0.95 [8]. In addition, there is considerable variation in CCT between different ethnic groups. Aghaian et al. [15] found that African-Americans (mean CCT of 524 µm) and Japanese (mean CCT of 538 μm) had significantly thinner corneas than the general population. Others [10] have not observed the same differences for Japanese populations; however; thinner corneas are consistently observed in the African-American populations [10, 13].

Genome-wide association studies (GWAS) on different human populations have identified a number of human loci/genes [16–18] associated with CCT. Several of the loci contain genes associated with CCT that are risk factors for human diseases, including *ZNF496*, which is associated with brittle cornea syndrome [19], and *COL8A2*, causing Fuch’s endothelial corneal dystrophy [20, 21]. A recent meta-analysis of over 20,000 individuals identified an additional 16 additional loci associated with CCT [22]. Six of these loci conferred a significant risk for keratoconus, a disease characterized by an extremely thin cornea. This study of a large population successfully identified many different genes that contribute to the heritability of CCT and not surprisingly implicated collagens and extracellular matrix pathways in the regulation of CCT. Although this represents a significant increase in the number of genes involved in CCT, all of the loci only account for approximately 8% of the additive variance of CCT in the European population [22]. Taken together, these data reveal that this highly heritable trait, CCT, is a complex trait. In fact, it could be so complex that the contribution of many individual genomic elements would be difficult to prove, especially in a genetically heterogeneous species like humans.

Attempting to define the link between CCT and POAG in a human GWAS is complicated by the fact that one is comparing two complex traits. In this case, the effect size has to be very large. Many of the early studies specifically looked for a genetic/molecular association between CCT and POAG. The large GWAS meta-analysis of human CCT [22] identified many new loci associated with CCT. One of the loci that conferred a significant risk for keratoconus (*FNDC3B*), was also associated with POAG. Given the power of the study (20,0000 individuals) and the small effect of individual SNPs on CCT, it is clear that the association of individual SNPs with CCT and POAG will require an extremely larger (potentially unrealistic) sample size. Leveraging the unique genotype of recombinant inbred mouse strains can help to simplify such analysis.

The approach taken in the present study was to identify genomic loci modulating CCT using the largest recombinant inbred mouse strain set, the BXD strain set. The BXD RI strains were produced by crossing the C57BL/6J mouse with the DBA/2J mouse. The progeny were inbred (brother-sister matings) to produce over 80 inbred strains. The first 42 of these strains are from the Taylor series of BXD strains generated at the Jackson Laboratory by Benjamin Taylor [23]. BXD43 and higher were bred by the Williams group at the University of Tennessee [24]. The BXD RI strains offer a powerful tool to accurately identify genomic loci modulating phenotypes such as CCT. Among the strain set, there are over 7,000 breakpoints in the genome. All of the strains are fully mapped and the parental strains are fully sequenced. Thus, SNPs as well as insertions or deletions are known for every of the BXD strains. With the aid of the bioinformatic tools present on GeneNetwork (Genenetwork.org), we were able to identify genomic loci and candidate genes modulating CCT in the mouse. We then used these data to interrogate human GWAS studies of corneal datasets [22] and glaucoma datasets [25, 26] to relate our findings in the mouse to the normal human cornea and disease states [27]. Ultimately, we examine genes that modify CCT in the mouse that may also be risk factors for human glaucoma.

## Results

The BXD RI strains were used to define genomic loci that modulate CCT. The CCT was measured in a total of 818 mice across 61 members of the BXD strain set (Figure 1). For the purpose of the present study we examined the total thickness of the cornea and no attempt was made to measure the separate layers of the cornea (epithelium, stroma or endothelium). The mean and standard error for each strain are shown in Figure 2. The mean central corneal thickness measured across 61 BXD strains was 100.9 μm with a standard deviation of 7.4 μm. The strain with the thinnest cornea was BXD 44 with an average corneal thickness of 85.3μm. The strain with the thickest cornea was BXD 74 with an average thickness of 124.6μm. The CCT for the parental strains is an intermediate value with the DBA2/J (D2) mouse having a CCT of 103.0μm and the C57BL/6J (B6) mouse having an average CCT of 93.1μm. Thus, there is considerable genetic transgression of CCT across the BXD RI strains with some BXD strains having corneas thinner than the parental strains and other strains having corneas thicker than the parental strains. This distribution of the CCT phenotype indicates that CCT is a complex trait, with multiple genomic loci segregating across the BXD RI strain set.

**Figure 1.**
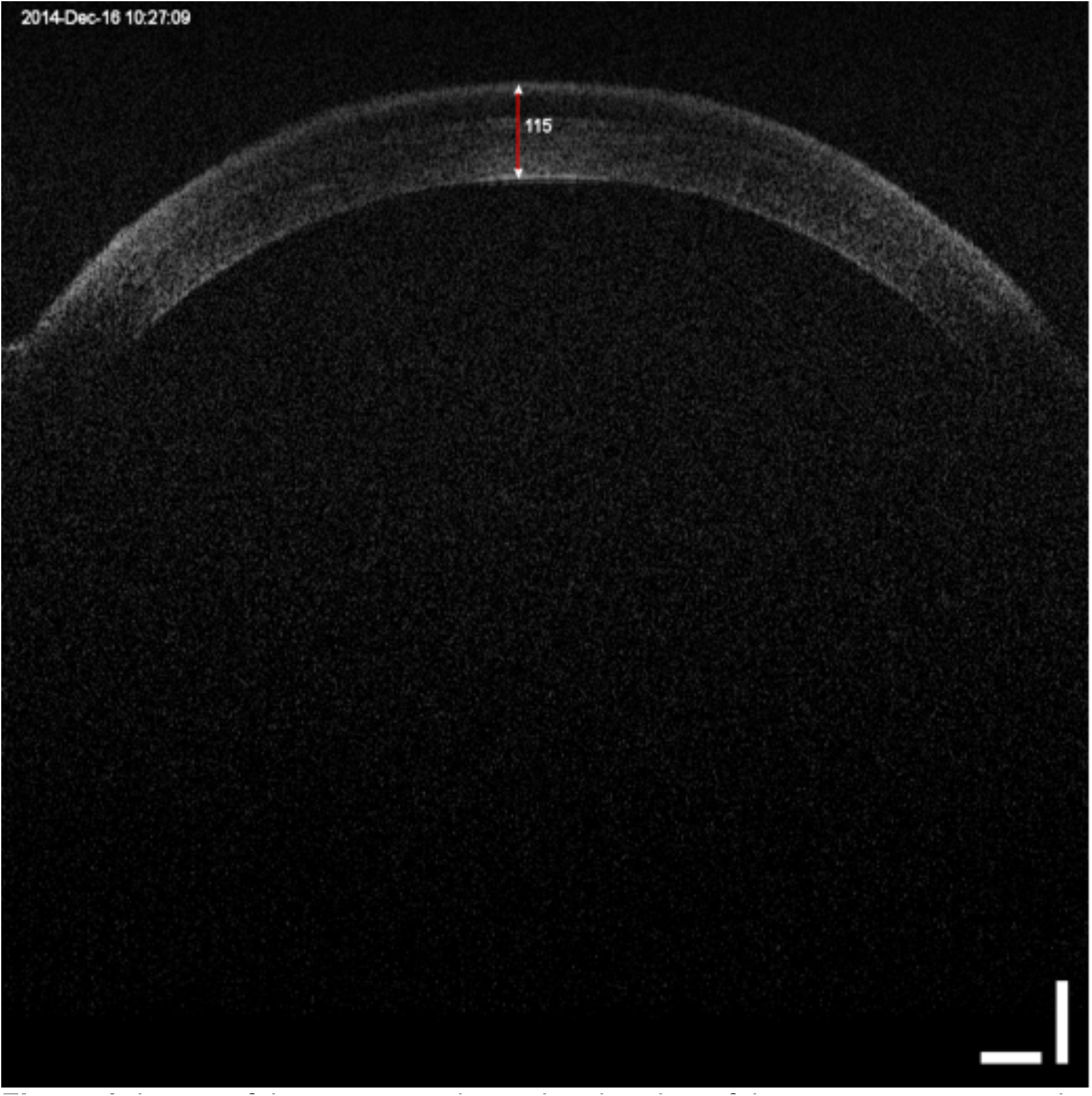
Image of the cornea and anterior chamber of the mouse eye scanned by the Phoenix Micron IV OCT System. The red bar shows the measurement of the central corneal thickness in microns.

**Figure 2.**
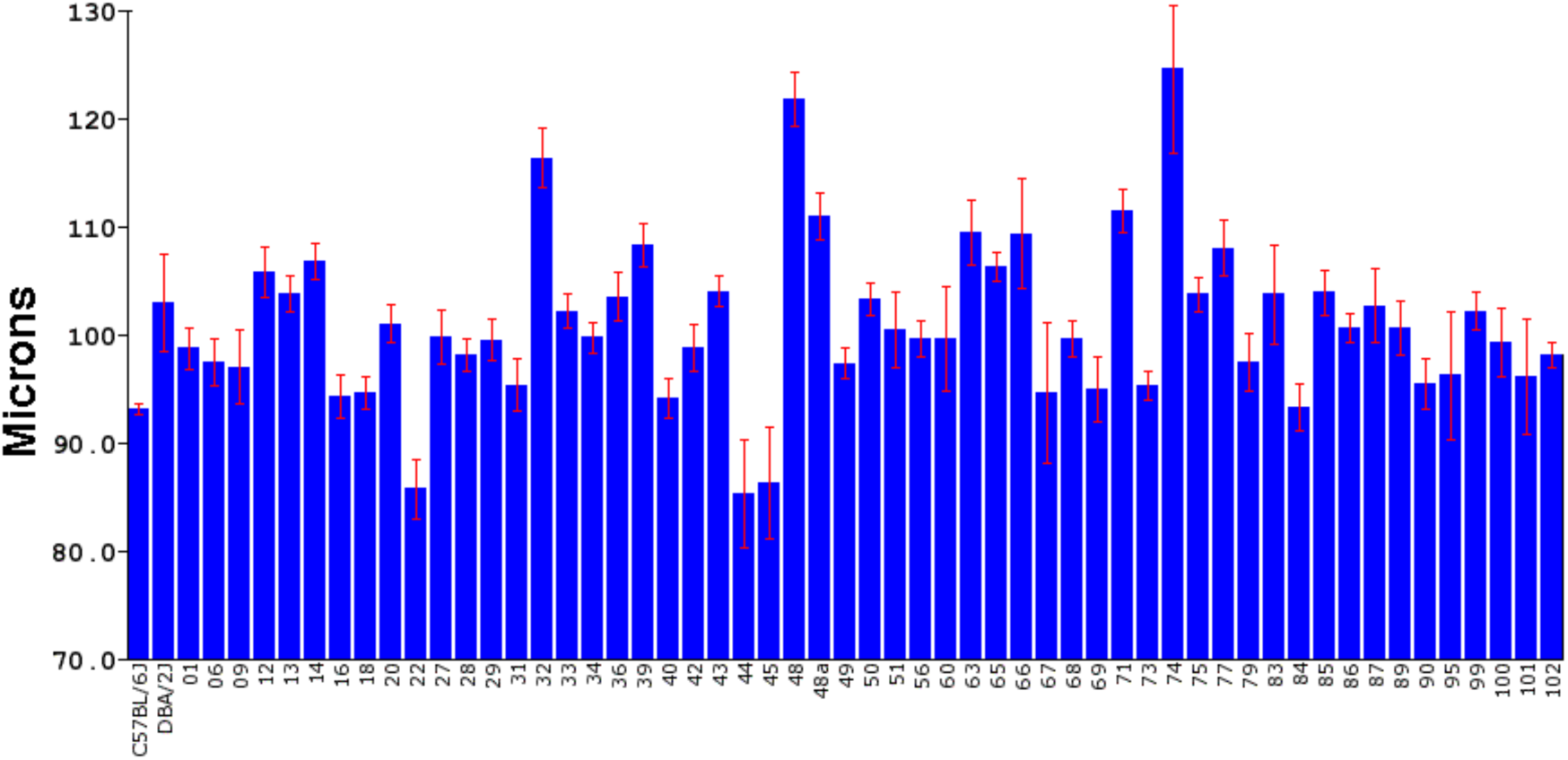
The CCT for 61 BXD strains is illustrated, with the mean indicated by the bar and standard error of the mean is presented by the red brackets. The specific BXD strains are indicated along the X-axis. The thickness of the cornea is shown to the right in microns.

These data also demonstrate that CCT is a heritable trait. Figure 2 shows that there is considerable variability in the CCT from strain to strain and the standard error for each strain is rather small, which suggests that the genetic variability has a greater effect than the environmental variability. These data can be used to calculate the heritability of CCT in the BXD RI strains. Heritability (H^2^) is the genetic variance (Vg) of the trait divided by the sum of genetic variance plus the environmental variance (Vg +Ve). The genetic variance can be estimated by calculating the standard deviation of the mean of the CCT for each strain (Vg = 10.054). The environmental variance can be estimated by taking the mean of the standard deviation across the strain (Ve = 2.92). Using the formula for heritability, H^2^ = Vg/(Vg + Ve), the calculation of 10.054/(10.054 + 2.92) reveals that H^2^ = 0.78. Thus, CCT is a highly heritable trait across the BXD RI strains.

Using an unbiased forward genetic approach and the data collected from the cohort of 61 BXD strains, we performed a genome-wide scan in an effort to identify QTLs that modulate CCT. The genome-wide interval map (Figure 3) reveals one significant peak on chromosome 13 (13 to 19 Mb) and a suggestive peak on chromosome 7 (42 to 57 Mb). An expanded view of the peak on chromosome 13 is illustrated in Figure 4 along with a haplotype map of the BXD strains. The strains with the thicker corneas in general tend to have the D2 allele; while, strains with thinner corneas have the B6 allele. This is reflected in the genome wide map by the presence of the green line in the peak on chromosome 13 that indicates that higher phenotypic values are associated with the D2 allele (Figure 3). The QTL on Chr. 13 (Figure 4) rises above the significance level of p<0.05 as indicated by the pink line. The significant portion of the peak extends from 13Mb to 19Mb over Chr. 13. Genomic elements modulating CCT are located in this region. To identify candidates for modulating CCT in the BXD RI strains, we examined this 7Mb long region. The candidate genes can either be genomic elements with cis-QTLs or they can be genes with nonsynonymous SNPs changing protein sequence. Within this region are 29 traditional genes and one microRNA. In the Whole Eye Database (Eye M430V2 (Sep08) RMA) hosted on GeneNetwork.org we found two genes within this locus with cis-QTLs: *Cdk13* and *Mplkip.* We examined the expression of both of these genes in the cornea and the eye using quantitative PCR (Supplemental data 1, S1 Appendix). Neither of the two genes were expressed at a high enough level in the cornea to be monitored by microarrays of the whole eye. Thus, both genes were discounted as potential candidates for modulating CCT. Within the significant QTL on Chr 13, only one gene (*Pou6f2)* had nonsynonymous SNPs. *Pou6f2* contained five separate nonsynonymous SNPs (rs29821949, rs52634762, wt37-13-18331131, rs29234524 and rs29250924). Three of these SNPs result in a change in the amino acid proline, which could cause a change in secondary structure of the protein and an alteration in protein function. Analyzing these changes using SIFT to predict functional changes in the protein structure [28], one SNP (rs29234524) was predicted to potentially alter the function of the protein. Based on this examination of the locus, there are no valid cis-QTLs and only one gene, *Pou6f2*, with nonsynonymous SNPs that could have biological effects.

**Figure 3.**
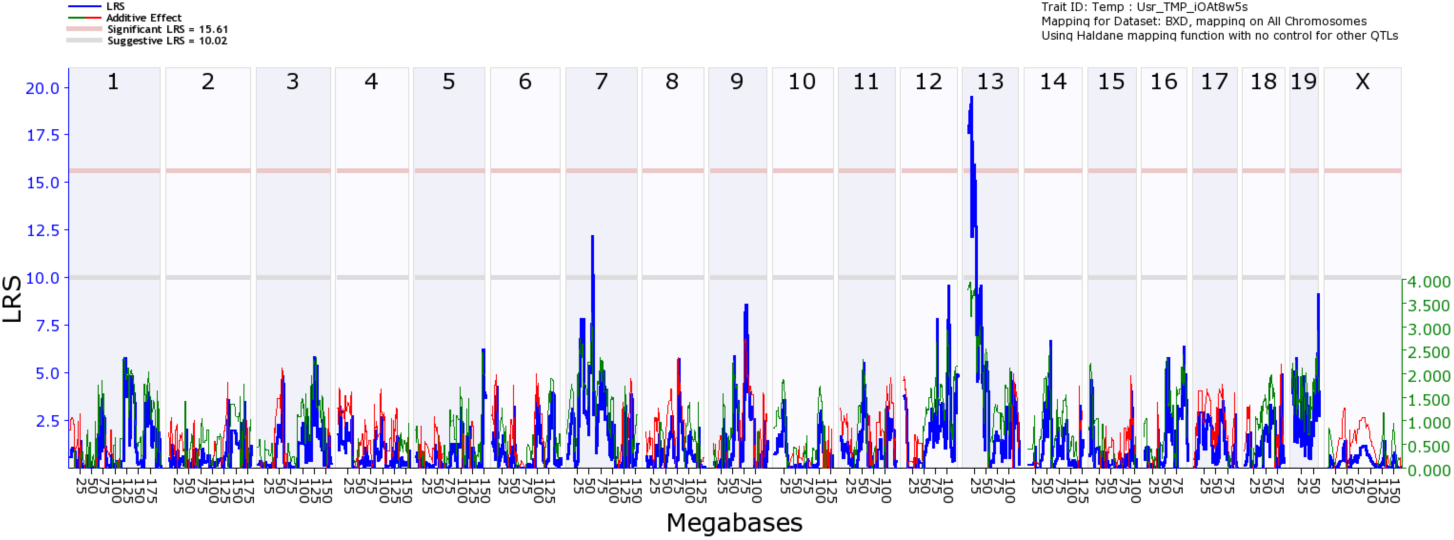
Interval map of CCT across the mouse genome is illustrated. The blue line indicates the total LRS score. The red line illustrates the contribution from the B6 allele and the green line the contribution from the D2 allele. The location across the genome is indicated on the top from chromosome 1 to chromosome X. On the y-axis is the linkage related score (LRS). Notice one significant QTL peak on Chr13 (above the pink Line, p = 0.05) and additional suggestive peaks (above the gray Line).

**Figure 4.**
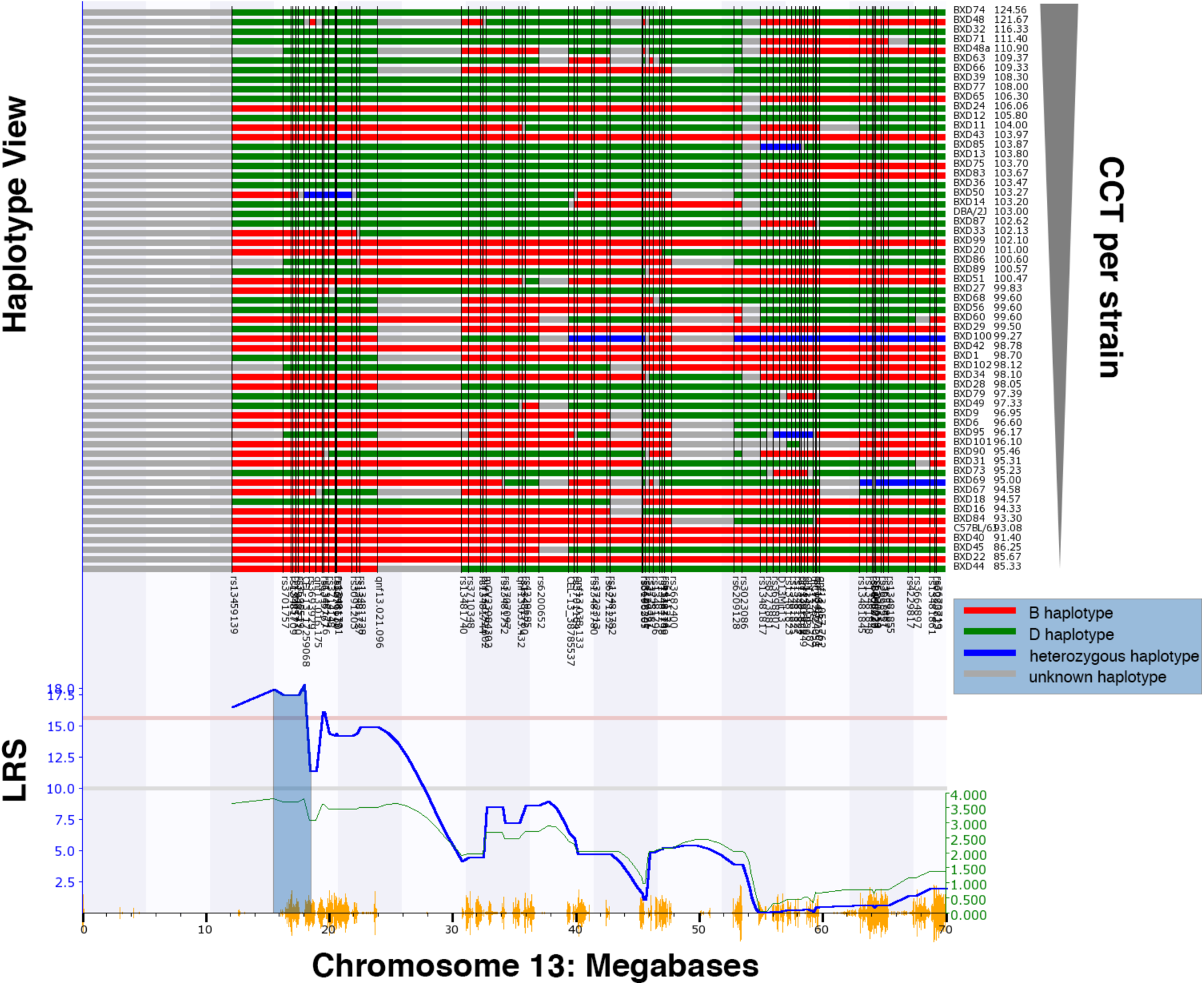
Map of gene locations across Chr. 13. The haplotype map for the 61 strains in the CCT dataset is shown in the top panel. As indicated at the key on the right, red represents the B6 alleles, green defines the D2 alleles, blue represents regions of the DNA that are heterozygotic and gray is unmapped. The genomic markers used in the mapping process are listed at the bottom of the haplotype map. At the far right is a list of the specific BXD RI strains and the associated CCT measurements going from thick corneas at the top to thin corneas at the bottom. QTL map of CCT on Chr. 13 is shown below the haplotype map. An LRS of above the pink line is statistically significant (p<0.05). A positive additive coefficient (green line) indicates that D2 alleles are associated with higher trait values. The light fill under the peak defines the genomic locus associated with the modulation of CCT. The significant peak for CCT is from 13 to 19 Mb (gray shaded area). Vertical orange lines at the bottom of the plot show the SNPs on Chr. 13.

The genomic elements within significant QTLs on mouse chromosome 13 (13 to 19 Mb) were examined to determine if there were specific associations in two human datasets: the International Glaucoma Genetics Consortium dataset for association with CCT [22], and the NEIGHBORHOOD glaucoma dataset [29]. The syntenic regions on human chromosome 1 (235 to 238 Mb) and chromosome 7 (38 to 43 Mb) were queried for associations of genetic effects relative to human CCT and with genetic risk for human glaucoma. In the International Glaucoma Genetics Consortium dataset for CCT [22], there were a number of nominally significant associations; however, none of the markers remained significant after corrections for multiple testing. When we looked specifically for the candidate genes from the mouse, weak associations were found in both European populations (peak marker rs4723833; P=4.34x10^−3^ β=-0.062µm) and Asian populations (peak marker rs17619647; P=3.72x10^−3^; β=-0.066µm), both of these SNPs (rs4723833 and rs17619647) overlap *POU6F2*. However, the associations are not significant after correction for multiple testing of 482 SNPs in Europeans and 265 SNPs in Asians. The β values from this linear regression analysis reflect the quantitative change in CCT per associated allele. The mouse locus was also used to interrogate the NEIGHBORHOOD human POAG glaucoma dataset [29]. In the NEIGHBORHOOD dataset, the top 50 SNPs associated with glaucoma within the syntenic region were all found within the *POU6F2* locus. The most significant SNP (rs76319873) had a genome-wide p-value of 5.34^−6^ and the next top 3 SNPs had p-values less than 10^−5^ (Figure 5, Table 1). None of these SNPs reached genome-wide significance when corrected for multiple testing.

**Table 1.** The four SNPs with the highest p values from the NEIGHBORHOOD Human GWAS Dataset. None of the SNPs reached genome-wide significance which is 6.4X10^−8^.

| CHR | BP | SNP | P | Gene |
| --- | --- | --- | --- | --- |
| 7 | 39318083 | rs76319873 | 5.34E-06 | POU6F2 |
| 7 | 39339109 | rs73130485 | 1.55E-05 | POU6F2 |
| 7 | 39333177 | rs75082513 | 3.52E-05 | POU6F2 |
| 7 | 39304125 | rs4302750 | 4.88E-05 | POU6F2 |

**Figure 5.**
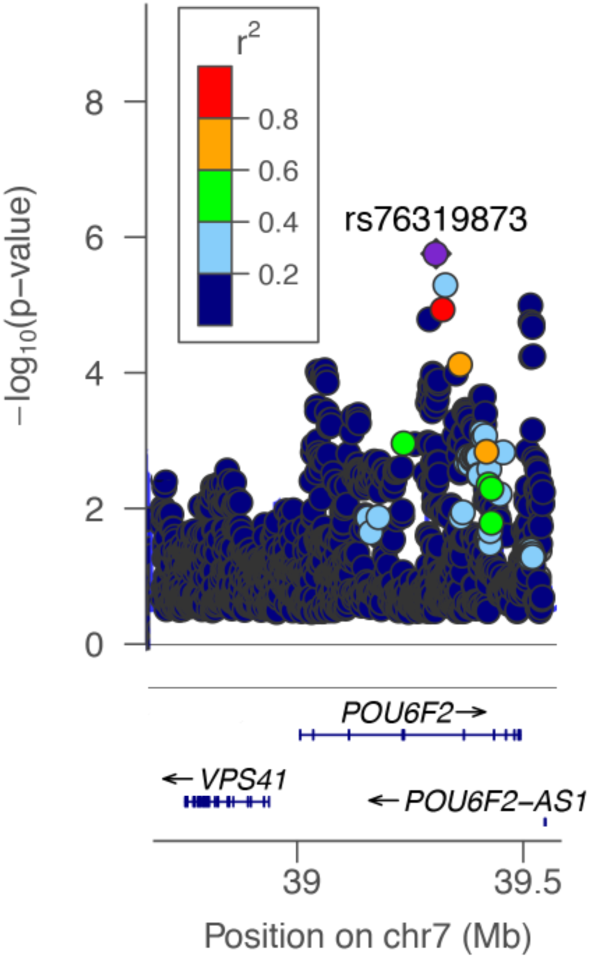
This is a Manhattan plot over the locus containing *POU6F2*. To the right is the p value related to glaucoma in a log_10_ scale. On the bottom portion of the panel the location of three human genes is shown: *VPA41, POU6F2* and *Pou6F2-AS1*. The SNP (rs76319873) with the highest association to POAG is colored purple. The panel insert denotes the imputed quality score (r^2^) and the appropriate SNPs are colored in the plot.

## Distribution of POU6F2 in the Mouse Eye

Our data suggests that *Pou6f2* modulates central corneal thickness and it may also modulate risk for glaucoma in humans. To determine if *Pou6f2* could be a molecular link between CCT and glaucoma, we examined the distribution of POU6F2 protein and mRNA in the retina and cornea in mice. The first approach was to examine its absolute expression levels in adult retina and cornea using digital droplet PCR. We first confirmed the specificity of tissue dissection using the corneal marker Uroplakin 1[30], which was highly abundant in cornea but not retina (5760±260 copies/µL vs. 3.3±0.9 copies/µL). The expression levels of *Pou6f2* were relatively high in the retina (76.3±1.5 copies/µL); however, levels of *Pou6f2* message from cornea were just above the detection threshold of digital droplet PCR (0.5±0.2 copies/µL), suggesting that only few if any corneal cells express *Pou6f2*. These data indicate that *Pou6f2* was barely expressed in the adult cornea, while it was readily detectable in the adult retina.

Previous studies [31] demonstrated that POU6F2 is expressed in neuroblasts in the future ganglion cell layer of the developing eye and in retinal ganglion cells in the adult mouse, cat and monkey. To confirm these findings and to examine the potential for a link between RGCs and cornea, we immunostained sections from the embryonic mouse eye, the adult mouse eye and flat-mounts of mouse retina. In the embryonic eye, there is strong staining of the neuroblasts destined to become retinal ganglion cells (Figure 6). There is also notable staining of the epithelium of the developing cornea and what appears to be corneal stem cells (Figure 6). The distribution of POU6F2 in the developing eye indicates a clear association between the development of retinal ganglion cells and the cornea.

**Figure 6.**
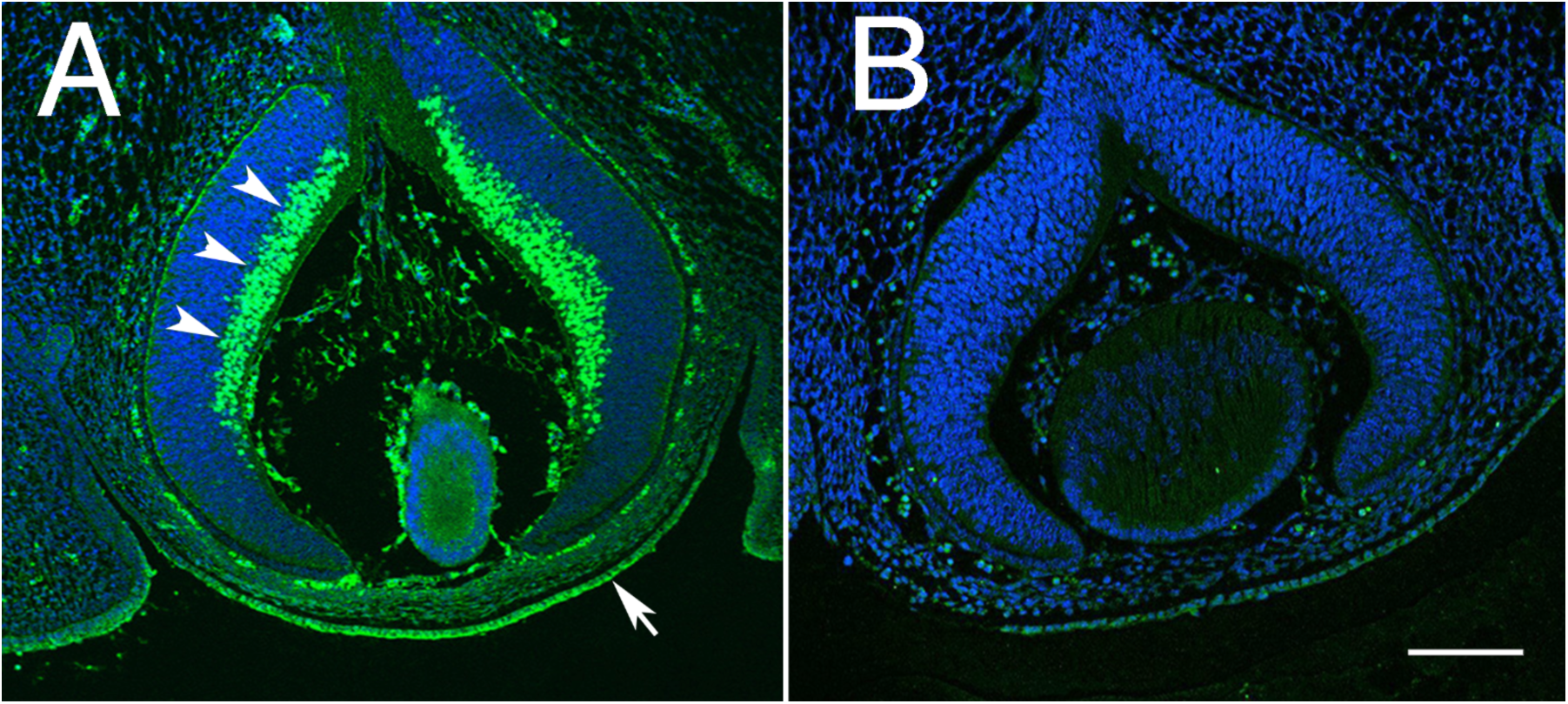
Distribution of POU6F2, in the embryonic eye P15 is illustrated. In sections stained for POU6F2 (A), there is prominent staining of neuroblasts destined to become retinal ganglion cells (arrow heads). There is also staining of the developing cornea and corneal epithelium (arrow). This staining is specific to the primary antibody for it is not present in sections in which the primary antibody was omitted (B) a secondary only control. Both sections are at the same magnification and the scale bar in B represents 100µm.

In sections through the adult mouse retina (C57BL/6J), POU6F2 is found to label the nuclei of many cells in the retinal ganglion cell layer. To fully explore the distribution of POU6F2 in the retinal ganglion cell layer, we examined flat mounts of the mouse retina (Figure 7) counterstained with RBPMS (a ganglion cell marker [32]) and a nuclear stain. We observed that POU6F2 labels only a small percentage of the total number of cells in the ganglion cell layer. When we quantified the number of cells labeled with POU6F2, only 17.4% of the total cells were heavily labeled with POU6F2. Virtually all of the POU6F2-positive cells were ganglion cells positive for RBPMS. There were a few cells in the inner nuclear layer that were positive for POU6F2 and these cells were also positive for RBPMS, identifying these cells as displaced ganglion cells. These data demonstrate that POU6F2 is expressed in a small subset of retinal ganglion cells. To provide additional evidence that the POU6F2 positive cells are ganglion cells, we crushed the optic nerve unilaterally in three mice and allowed them to survive for 28 days. The retinas were then examined for POU6F2 staining. No labeling was observed in the RGC layer, indicating that all POU6F2 cells were gone (Figure 1, S1 Appendix). Taken together, these data reveal that POU6F2 is expressed in a subset of retinal ganglion cells in the ganglion cell layer. Finally, as previously observed by Zho et al. [31] there were a few cells labeled with POU6F2 on the inner surface of the inner nuclear layer. These cells appeared to be displaced ganglion cells (Figure 2, S1 Appendix). Thus, POU62 labels a subset of RGCs within the mouse retina.

**Figure 7.**
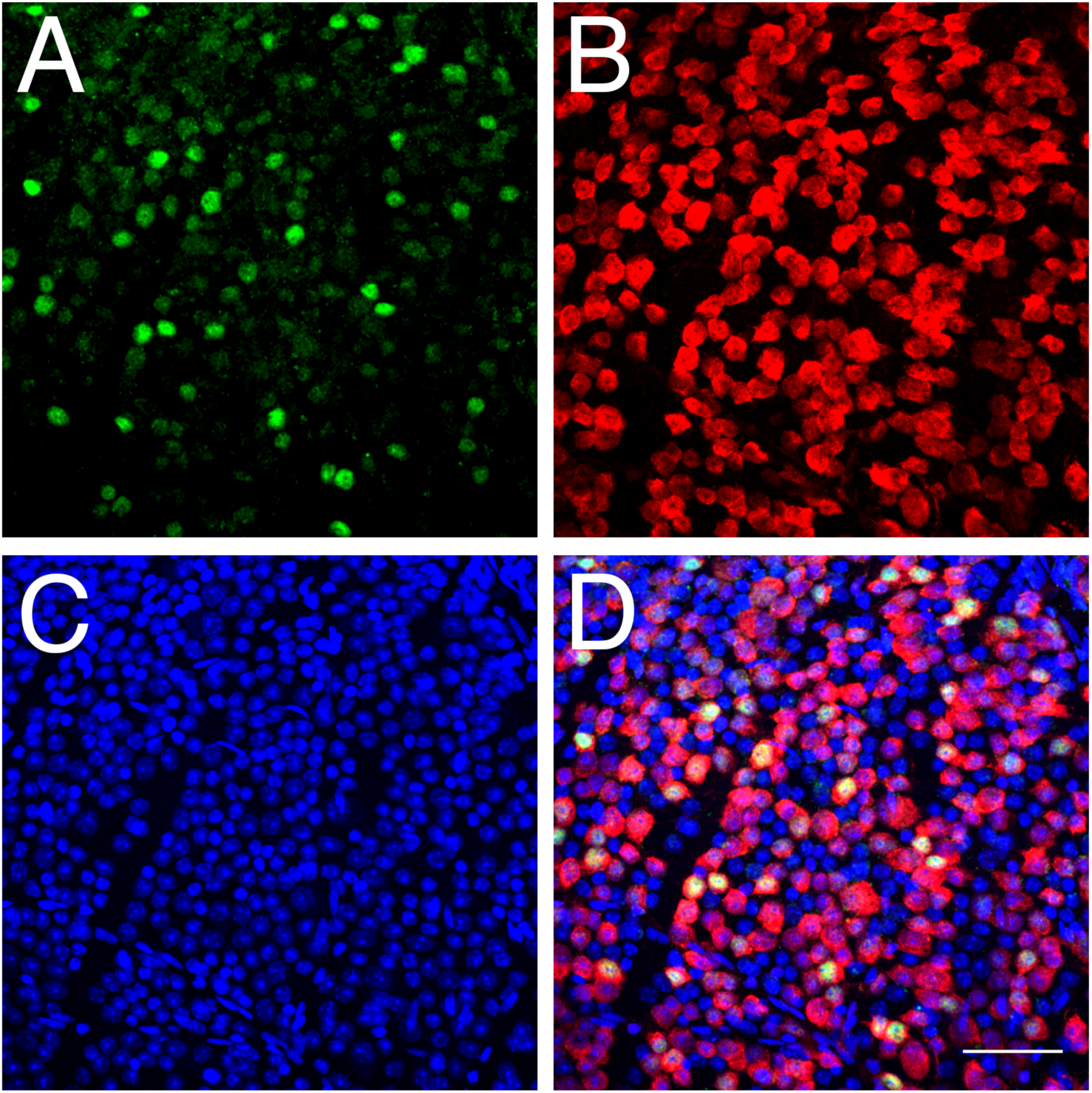
The labeling of RGCs with an antibody directed against POU6F2 is illustrated in a flat-mount of the mouse retina. Panel A is the staining within the retina (green) for POU6F2. The same area (panel B) was stained (red) for the ganglion cell marker RBPMS. Nuclei in the ganglion cell layer were stained blue with TOPRO-3 (panel C). The merged image is shown in D. Notice that many of the ganglion cells were heavily stained for POU6F2 and that others are lightly labeled. The scale bar in D represents 50µm.

To determine if the POU6F2-positive RGCs are selectively sensitive to glaucoma, we double stained 4 young DBA/2J (2 months old) mice for POU6F2 and RBPMS and compared these values to 4 aged (8-month-old) DBA/2J mice (Figure 8). When we counted the cells in the 4 young mice there were an average of 460 (±81, SEM) RBPMS-labeled RGCs per 40X field. In the same sections, we observed an average of 74 POU6F2-labeled RGCs (±15). When these results were compared to 4 aged DBA/2J mice (8 months old) there was a significant decrease (p<0.001, student t test) in the total number of RGCs to an average of 361 RGCs (±21) per 40 X field. This represented a total decrease in RBPMS labeled ganglion cells of 22%. When we examined POU6F2-positive cells in the aged DBA/2J retinas there was a 73% loss of loss of cells, a significant difference relative to young DBA/2J retinas (p<0.0001). The average number of POU6F2 cells in the young retina was 74 (±5), while in the aged retina this number decreased to 21 (±3). These data demonstrate that the POU6F2 RGC subtype is very sensitive to early phases of glaucoma in the DBA/2J model.

**Figure 8.**
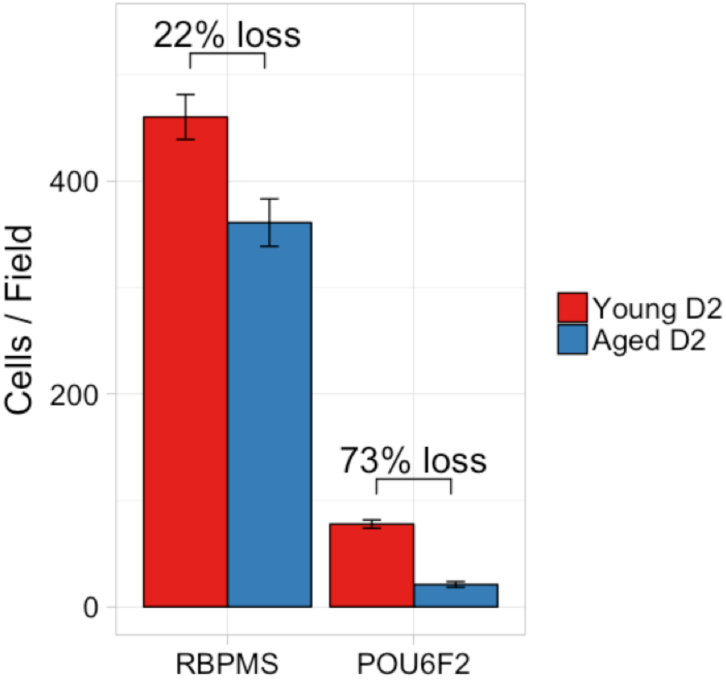
The selective sensitivity of POU6F2 RGC subtypes is demonstrated using the DBA/2J model of glaucoma. There was a 22% loss of RBPMS-labeled RGCs in aged DBA/2J mice (8 months of age, Aged D2) as compared to young DBA/2J mice (2 months of age, Young D2). There was a dramatic loss of 73% of the POU6F2-positive cells comparing the Young D2 mice to the Aged D2 mice. These data demonstrate the sensitivity of the POU6F2 RGC subtype to glaucoma.

To examine the expression of POU6F2 in the cornea, we first examined sections through the cornea and limbus extending down into the sclera. There was no staining in the cornea or sclera. Occasionally, we observed individual cells stained in the limbal area. However, it was difficult to unequivocally identify these cells as limbal stem cells. As an alternative approach, we stained the surface of six eyes using extended antibody incubation times. When examining the junction between the cornea and sclera, a line of cells is stained at the junction between the cornea and sclera in the limbus (Figure 9). This line of cells was also positive for the stem cell marker ABCB5 (Figure 9E). Thus, POU6F2 is not only a marker for limbal stem cells but it may be responsible in part for the maintenance of the corneal integrity in the adult mouse eye.

**Figure 9.**
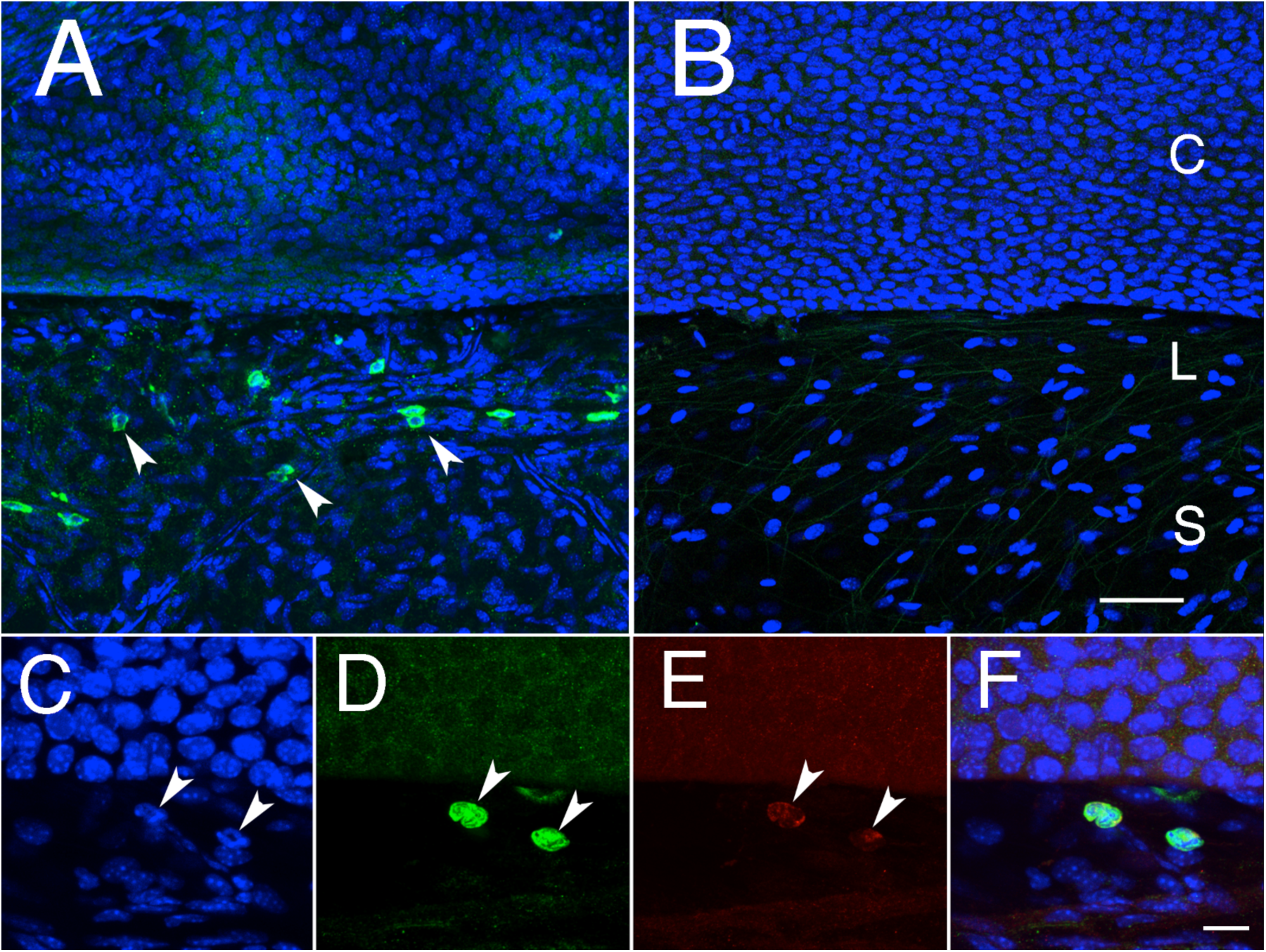
The surface of the eye was stained for POU6F2 and counter-stained with TOPRO-3 (A). The control section without primary antibody is shown in B. The limbus (L) is the transition between the cornea (C) and the sclera (S). In A, there are many cells stained for POU6F2 and all of these cells are localized to the limbus. C, D, E and F are higher magnification photomicrographs taken from the same region of a triple stained section. The section was stained with TOPRO-3, a nuclear marker (C), POU6F2 (D) and ABCB5 (E). F represents a merged image of the three channels. Two stem cells are labeled by arrows in C, D and E. Photomicrographs in A and B are taken at the same magnification and the scale bar in B represents 50µm. Photomicrographs in C, D, E and F are taken at the same magnification and the scale bar in F represents 20 µm.

To determine if POU6F2 is directly involved in corneal development and maintenance affecting CCT, we examined the corneas of 8 *Pou6f2*-null mice and compared their CCT to that from 10 wild-type littermates (Figure 10). There was a significant difference (p<0.01, Wilcoxon signed-rank test) in CCT between the *Pou6f2*-null mice and the wild-type littermates. In the *Pou6f2*-null mice the mean CCT was 102.8µm (±1.8µm). The wild-type littermates had a mean CCT of 109.9µm (±1.8µm). Thus, abolishing *Pou6f2* expression has a significant effect on CCT in the adult mouse.

**Figure 10.**
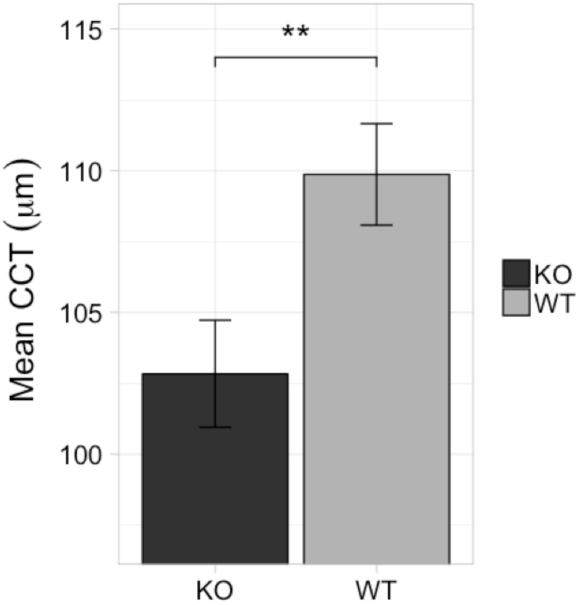
The effect of knocking out *Pou6f2* on CCT. The average CCT (bars represent SEM) from Wild-type (WT) mice and *Pou6f2*-null (KO) mice is shown. In the Wild-type littermates (n=10) the cornea was on average significantly (p < 0.01) thicker than observed in the *Pou6f2*-null mice (n=6).

## Discussion

The BXD RI strain set is particularly well suited for a systems genetic approach to study CCT. This relatively new branch of quantitative genetics has the goal of understanding networks of interactions across multiple levels that link DNA variation to phenotype [33]. The present approach involves an analysis of sets of causal interactions among classic traits such as CCT, together with networks of gene variants, and developmental/environmental factors. The main challenge is the need for a comparatively large sample size and the use of more advanced statistical and computational methods and models. The BXD strain set is sufficiently large to have adequate power for this approach [34, 35]. The genetic reference panel used in this study consisted of 62 strains of mice. The novel aspect of our current approach is the combination of phenotypic mouse data, mRNA expression data (baseline and experimental) from the same BXD RI strains and the interrogation of GWAS studies for glaucoma (NEIGHBORHOOD Study [25]) and for CCT [22]. Data that we generated throughout this experiment and data already deposited in GeneNetwork (<u>www.genenetwork.org</u>) for gene expression in the eye [34] and retina [36–38] were used to test specific mechanisms and predictions.

In the present study, we have identified a significant QTL on Chr 13 (13-19 Mb) that modulates CCT in the BXD RI strain set. One suggestive QTL was also found on chromosome 7 (41 to 57 Mb). Previous studies from the Anderson group [39, 40] have found significant QTLs associated with CCT. Using a F2 cross between C57BLKS/J and SL/J F2 mice, Liverly et al. [39] found a significant QTL on Chr. 7 at 105Mb that modulated CCT. This differs from the suggestive QTL identified on Chr. 7 in the present study. The QTL identified with the C57BLKS/J and SL/J F2 cross is located at 105Mb on Chr7, while the suggestive QTL from the present study was at 41 to 57Mb on Chr7. A second study by Koehn et al. [40], identified two significant QTLs associated with CCT in an F2 cross between BXD24/TyJ and CAST/EiJ. One was on Chr 3 at 104Mb, and the second was on Chr11 at 88Mb [40]. Neither of these loci demonstrated even suggestive associations with CCT in the BXD RI strains used in the present study.

In humans, CCT is a highly heritable trait affected by many genomic loci [16–22]. Comparing these human loci to those identified in the mouse reveals several genomic regions that are similar. For example, the mouse locus on Chr7 at 105Mb [39] could include previously identified human loci near several known genes: TJP1, CHSY1, LRRK1 and AKAP13 [22, 41]. Other than this region in the mouse, there does not appear to be any overlap between loci modulating CCT in the mouse and those identified in the human.

CCT is a phenotypic risk factor for glaucoma [1–3]. Glaucoma is a diverse set of diseases with heterogeneous phenotypic presentations associated with different risk factors. Untreated, glaucoma leads to permanent damage of axons in the optic nerve and visual field loss. Millions of people worldwide are affected [42, 43] and glaucoma is the second leading cause of blindness in the United States [44]. Adult-onset glaucoma is a complex collection of diseases. The severity of glaucoma appears to be dependent on the interaction of multiple gene variants, age, and environmental factors [45]. These complex genetic risk factors are the focus of recent genome-wide association studies that are not only defining genetic risk factors but also aiding in our understanding of disease mechanisms [46–48]. These would include the identification of genes that could influence cerebral spinal fluid pressure [49] and ultimately the differential pressure observed across the optic nerve head [50]. Recent findings suggest that RGC number is also a risk factor for glaucoma. Several polymorphisms in the genes *SIX6* and *ATOH7* were identified that are responsible for thinner retinal nerve fiber layer due to fewer RGCs, suggesting that humans with fewer RGCs are at increased risk of developing glaucoma [26, 51, 52]. In addition, linkage analysis in families affected by glaucoma has led to an extensive list of genomic loci linked to POAG. The genes underlying some of the adult-onset glaucoma loci have been identified (*MYOC*, *OPTN*, *TBK1, CDKN2BAS* [53] [54] [55, 56].

Many studies in human and mouse have looked for a genetic link between CCT and glaucoma. Studies in the human cornea have found several SNPs that modulate corneal thickness. However, none of the genes in these studies appear to relate CCT to the risk of developing glaucoma with the exception of *FNDC3B*[22]. Several studies have examined the genomic loci in the mouse that modulate CCT and again none of the loci controlling CCT thickness relate to glaucoma risk. In the present study, we have identified a well-defined QTL associated with CCT in the mouse. Within this locus, there is a limited number of candidate genes. One gene, *Pou6f2,* is uniquely poised to modulate CCT, for we have shown that it is expressed in the developing cornea and in limbal stem cells. In addition, *Pou6f2* may also be associated with glaucoma risk. Interestingly, Zhou et al. [31] demonstrated that *Pou6f2 i*s present in the developing and adult retinal ganglion cells. This gene product is also associated with the regeneration of neurons in zebrafish [57].

Based on the distribution of POU6F2 in the cornea and retinal ganglion cells in the developing and adult mouse eye, it is tempting to suggest that *Pou6f2* may be a molecular link between CCT and the potential for RGC loss in glaucoma. A search of the normal mouse retina (DoD Normal Retina Affy MoGene 2.0 ST [37]) on GeneNetwork (genenetwork.org) reveals that *Pou6f2* forms a genetic network with retinal ganglion cells markers, including *Thy1, Pou4f1* (Brn3a) and *Rbfox3* (NeuN) [58]. This association suggests that *Pou6f2* could be commonly regulated in a specific subset of retinal ganglion cells. These RGC subtypes may be selectively sensitive to loss in mouse models of glaucoma. In the present study, we have also shown that *Pou6f2* is expressed in the developing cornea and corneal stem cells. When we examined the NEIGHBORHOOD database, *POU6F2* has an interesting association with POAG. However, for the International Glaucoma Genetics Consortium dataset for CCT [22] the P value was p = 0.0037, and when corrected for multiple testing it did not reach a significant level. Thus, we are not able to definitively say that *Pou6F2* is a molecular link between CCT and glaucoma. As discussed above, CCT is a highly heritable trait and it is the second highest risk factor for glaucoma, following IOP. If this is the case, then why do we not know the molecular link between CCT and POAG? Some phenotypic traits are very complex or even hyper-complex. A well-known example of a complex trait is height. It is estimated that there are over 2,000 genomic loci that may contribute to height and that the heritability is relatively high as much as 80% [59], although others estimate that the heritability is lower due to dependence on family-based studies indicating that the true heritability is between 60% to 70% [60]. Nonetheless, the weight of any one genomic locus influencing height in the human population is low and genetic components known to be associated with heritability can only be accounted for a portion of the heritability. That being said, CCT is a glaucoma risk factor and a highly heritable trait. The identification of molecular links connecting CCT to POAG may be complicated by the fact that both are complex traits, influenced by multiple genomic as well as environmental factors. Using a mouse model like the BXD RI strain set (a restricted genetic reference panel) is an effective approach to identify genomic elements modulating phenotypes such as CCT. These data can now be used to design experiments that will attempt to link CCT to potential glaucoma risk.

POU6F2 appears to mark a subpopulation of RGCs that are selectively sensitive to injury in the DBA/2J mouse model of glaucoma. To understand the potential role of POU6F2 we examined data from a ChIP-chip experiment [61] in HEK293 cells. In this human cell line, POU6F2 bound to a number of different targets: CHD5, DACH1, ELAVL3, GFRA2, GRIN1, LHX1, MTSS1, NAP1L3, NEFH, NPTX1, NR4A2, NTNG2, PCDH7, ROBO2, SLC30A3, SLC7A8, SORCS2, UNC5A and VGF. If we examine these targets in the DoD Normal Retina Database [37] on GeneNetwork, all of the targets are expressed at relatively high levels in the retina, at least 2-fold above the mean expression of mRNA. In addition, their expression levels are highly correlated across the BXD strain set, with most probes having a Pearson’s *r* value above 0.7. One of these down-stream targets, *Slc30a3,* encodes a zinc transporter (ZNT3). Recent studies from the Benowitz lab [62], reveal that ZNT3 plays an important role in injured ganglion cells and axon regeneration. Thus, it is tempting to hypothesize that POU6F2 exerts its effects on retinal ganglion cell survival in an injury and chronic neurodegeneration (glaucoma) context by modulation of Slc30a3. In conclusion, we have shown that POU6F2 is involved in corneal development and modulates CCT. In retinal ganglion cells, POU6F2 is a transcription factor marking a subtype that is selectively sensitive to injury. POU6F2 is also known to be upstream of genes that play a critical role in ganglion cell death following injury and it is a potential glaucoma risk factor in humans.

## Materials and Methods

### Mice

All of the procedures involving mice were approved by IACUC at Emory University and the University of Tennessee Health Science Center. The study adhered to the ARVO Statement for the Use of Animals in Research. All of the mice in this study were between 60 to 110 days of age, which is long before any significant elevation in IOP due to two gene mutations (*Tyrp1* and *Gpnmb*) carried by selected BXD strains originating from the DBA/2J mouse. We examined equivalent numbers of males and females for each of the strains. Mice were housed in a pathogen-free facility at UTHSC or at Emory University, maintained on a 12 hr light/dark cycle, and provided with food and water ad libitum. All CCT measurements on the *Pou6f2*-null and wild-type mice were performed in a double-blind manner. The mice were genotyped after all CCT measurements were made.

### Measuring CCT

CCT was measured using two different Ocular Coherence Tomography (OCT) systems: a Bioptigen SD-OCT system (Morrisville, NC) and a Phoenix Micron IV OCT Imaging system (Pleasanton, CA). Mice were anesthetized using ketamine 100mg/kg and xylazine 15mg/kg. The eye was positioned in front of the lens. The entire anterior chamber was imaged. The corneal scans were saved to a portable hard drive for subsequent analysis. CCT was then measured three times for each eye using the Mouse Retina Program, InVivoVue Clinic, in the Bioptigen Software or the Micron OCT software. These data were stored and entered into an Excel spreadsheet. Repetitive measures were possible with the BXD RI strains since each strain has the identical genetic background allowing us to sample the developmental consequences of the same genetic background for all independent measures. For the BXD strains the corneas were measured between 60 and 90 days. In the *Pou6f2*-null mice the corneal thickness was measured at 30 days of age.

### Optic Nerve Crush

Optic nerve crush was performed as described in Templeton and Geisert [63]. Briefly, three C57BL/6J mice were anesthetized using ketamine 100mg/kg and xylazine 15mg/kg. Under the binocular operating scope a small incision was made in the conjunctiva. With micro-forceps (Dumont #5/45 Forceps, Roboz, cat. #RS-5005, Gaithersburg, MD), the edge of the conjunctiva was grasped next to the globe. The globe was rotated nasally to allow visualization of the posterior aspect of the globe and optic nerve. The exposed optic nerve is then clamped 2 mm from the optic nerve head with Dumont #N7 self-closing forceps (Roboz, cat. #RS-5027) for 10 seconds. At the end of the procedure, a drop of 0.5% proparacaine hydrochloride ophthalmic solution (Falcon Pharmaceuticals, Fort Worth TX) was administered for pain control and a small amount of surgical lube (KY Jelly, McNeil-PPC, Skillman NJ) was applied to the eye to protect it from drying. The mouse was allowed to wake up on a heating pad and monitored until fully recovered.

### Interval mapping for the traditional phenotypes

CCT data was subjected to conventional QTL analysis using simple and composite interval mapping and pair-scans for epistatic interactions. Genotypes were regressed against each trait using the Haley-Knott equations implemented in the WebQTL module of GeneNetwork [64, 65]. Empirical significance thresholds of linkage were determined by permutations [66]. We correlated phenotypes with expression data for whole eye and retina generated by Geisert, Lu and colleagues [34, 37]. Based on recent work with the enlarged set of BXDs, we expected all significant (p < 0.05) QTLs to be mapped with a precision of ± 2 Mb [35, 67–69]. To identify loci, and also to nominate candidate genes, we used the following approaches: interval mapping for the traditional phenotypes, candidate gene selection within the QTL region, cis-eQTL analysis of gene expression, trans-eQTL analysis, multi-trait and complex analysis of molecular, clinical, and laboratory traits.

### NEIGHBORHOOD Analysis

The peak area of association on mouse chromosome 13 was examined for associations in human datasets. Syntenic regions on human chromosomes 1 and 7 were queried for associations with POAG in the NEIGHBORHOOD [25] dataset. Subsequently, the POU6F2 locus was queried in the International Glaucoma Genetics Consortium dataset for association with CCT [22].

### RNA isolation and digital PCR

In order to quantify Pou6f2 mRNA expression, C57BL/6J mice (n=5) were deeply anesthetized with a mixture of 15 mg/kg of xylazine and 100 mg/kg of ketamine and sacrificed at 9am. Corneas and retinas were dissected separately and stored in Hank’s Balanced Salt Solution and RiboLock (Thermo Scientific, Waltham MA) at −80°C. RNA was isolated on a Qiacube with the RNeasy mini kit (both Qiagen, Hilden, Germany) according to the manufacturer’s instructions with additional on-column DNase1 treatment. RNA integrity was assessed using an Agilent Bioanalyzer 2100 and RIN scores for both pooled tissues were >9.5. Takara PrimeScript (Clontech, Mountain View, CA) was used to retrotranscribe equal amounts of RNA for both tissues. Digital Droplet PCR was then carried out using 30ng of total RNA in 20µL reactions of EvaGreen ddPCR supermix supplying 2µM Mg^2+^. Primers were designed using NCBI PrimerBlast to work at a combined annealing/extension temperature step of 60°C and the sequences were as follows: Upk1b fwd (5-CAGGCAGCCGGTCTTTTAGAAA-3), Upk1b rev (5-ATCATTGTTGGTGGCTTCGAGA-3), Pou6f2 fwd (5-CCCTCAATCAGCCAATCCTCAT-3), Pou6f2 rev (5-GTTCAGGGATGAGGTAGCTTGT-3). A combination of Ppia and Gapdh (Qiagen Quantitect primer assays) was used for ddPCR normalization.

### Immunohistochemistry

Twelve C57BL/6J mice were deeply anesthetized with a mixture of 15 mg/kg of xylazine and 100 mg/kg of ketamine and perfused through the heart with saline followed by 4% paraformaldehyde in phosphate buffer (pH 7.3). The eyes were removed and embedded in paraffin and sectioned at 5*µ*m. For the embryonic eyes, two timed-pregnant female mice were deeply anesthetized with tribromoethanol, decapitated and the E16 embryos were removed. The embryo heads were placed in 4% paraformaldehyde on ice for 1 hour, rinsed with PBS and dehydrated in stages of ethanol followed by xylenes, and then embedded in paraffin. The 5*µ*m sections of retina and E16 mice had the paraffin removed and the sections were blocked with 5% normal donkey serum and stained with a rabbit antiserum directed against POU6F2 (MyBiosource, Cat. # MBS9402684) at 1:500 for 2 hours at room temperature (Supplemental Data 2, S1 Appendix). To demonstrate the specificity of the POU6F2 antibody, retinas from *Pou6f2*-null mice were stained and no nuclear labeling was observed. The sections were rinsed three times and transferred to secondary antibodies (Alexa Fluor 488 AffiniPure Donkey Anti-Rabbit, Jackson Immunoresearch, Cat. #715-545-152; at 1:1000 for two hours at room temperature. After three washes of 15 minutes each, To-PRO-3 Iodide (Molecular Probes, Cat. # T3605) was applied 1:1000 as a nuclear counterstain. After a final wash in PBS, coverslips were placed over the sections using Fuoromount -G (Southern Biotech, Cat. #0100-01) as a mounting medium. For the retina flat mounts, the retinas were removed from the globe and rinsed in PBS, blocked in 5% normal donkey serum and placed in primary antibodies in primary rabbit antibody POU6F2 (MyBiosource, Cat. # MBS9402684) at 1:500 and to label RGCs, a primary guinea pig antiserum against RBPMS (Millipore, Cat # ABN1376) at 1:1000 overnight. The retinas were rinsed and placed in secondary antibodies, (Alexa Fluor 488 AffiniPure Donkey Anti-Rabbit, Jackson Immunoresearch, Cat. #715-545-152 and Alexa Fluor 594 AffiniPure Donkey Anti-Guinea pig, Jackson Immunoresearch, Cat. #706-585-148) at 1:1000 for two hours at room temperature. After three washes of 15 minutes each, To-PRO-3 Iodide (Molecular Probes, Cat. # T3605) was applied 1:1000 as a nuclear counterstain. After To-PRO3 has stained the section for 30 minutes, slides were washed twice with PBS for 10 minutes followed by a 5-minute wash with distilled water. Autofluorescence was decreased by treating the sections with cupric sulfate [70]. The 10mM cupric sulfate pH5 was applied to the section for 5 minutes. Slides were washed for 5 minutes with distilled water, then twice with PBS for 10 minutes and the slides were cover-slipped.

### Staining the Surface of the Limbus

Adult C57BL/6J mice were anesthetized with a mixture of 15 mg/kg of xylazine and 100 mg/kg of ketamine and perfused with saline followed by 4% paraformaldehyde in phosphate buffer (pH 7.4). The eyes were removed and post-fixed in 4% PFA for one at hour room temperature. The sclera along the ministry equator line was removed along with the posterior sclera and whole retina, vitreous body and lens. The remaining portion of the eye was rinsed three times in PBS containing 2% Triton X-100, 10 minutes each. Dissected blocks the anterior hemisphere were placed in 10% donkey serum and 4% BSA overnight 4 °C. The tissue was incubated for two days in rabbit primary antiserum to POU6F2 (MBS9402684 MyBiosource, San Diego CA) and ABCB5 (Product ab126 Abcam, Cambridge MA) at 1:500 at 4°C. The dissected limbal area was washed 3 times with PBS. Finally, the tissue was placed in secondary antibodies that included Donkey anti-Rabbit labeled with Alexa 488 (#711-545-152, Jackson ImmunoResearch Laboratories, West Gove CA) and Donkey anti-Goat IgG labeled with Alexa 594 (#705-585, Jackson ImmunoResearch Laboratories, West Gove CA) at 4 °C overnight. The tissue was cut into 5 pieces, radially mounted on a glass slide and a cover slip was placed over the tissue.

## Acknowledgements

We would like to thank XiangDi Wang (Department of Ophthalmology, Hamilton Eye Institute) for her assistance in collecting data. Department of Ophthalmology, Hamilton Eye Institute) for her assistance in collecting data. We would like to thank Micah Chrenek for breeding and genotyping all of the *Pou6f2*-null mice. Chelsey Faircloth was responsible for the sectioning and staining of the adult and embryonic eyes. Xue Jun Qin Duke Molecular Physiology Institute, Duke University, Durham NC.

